# Global quantitative analysis of the human brain proteome and phosphoproteome in Alzheimer’s disease

**DOI:** 10.1101/2020.05.19.105197

**Authors:** Lingyan Ping, Sean R. Kundinger, Duc M. Duong, Luming Yin, Marla Gearing, James J. Lah, Allan I. Levey, Nicholas T. Seyfried

## Abstract

Alzheimer’s disease (AD) is characterized by an early, asymptomatic phase (AsymAD) in which individuals exhibit amyloid-beta (Aβ) plaque accumulation in the absence of clinically detectable cognitive decline. Here we report an unbiased multiplex quantitative proteomic and phosphoproteomic analysis using tandem mass tag (TMT) isobaric labeling of human post-mortem cortex (*n*=27) across pathology-free controls, AsymAD and symptomatic AD individuals. With off-line high-pH fractionation and liquid chromatography coupled to tandem mass spectrometry (LC-MS/MS) on an Orbitrap Lumos mass spectrometer, we identified 11,378 protein groups across three TMT 11-plex batches. Immobilized metal affinity chromatography (IMAC) was used to enrich for phosphopeptides from the same TMT-labeled cases and 51,736 phosphopeptides were identified. Of these, 48,992 were quantified by TMT reporter ions representing 33,652 unique phosphosites. Two reference standards in each TMT 11-plex were included to assess intra- and inter-batch variance at the protein and peptide level. This comprehensive human brain proteome and phosphoproteome dataset will serve as a valuable resource for the identification of biochemical, cellular and signaling pathways altered during AD progression.

## Background & Summary

Alzheimer’s disease (AD) is the most common age-related neurodegenerative disease, currently affecting more than 47 million people worldwide^1,2^. AD is characterized by an early, asymptomatic phase (AsymAD) in which individuals exhibit amyloid-beta (Aβ) plaque accumulation in the absence of significant tau neurofibrillary tangles (NFT) and cognitive decline^3,4^. Currently the downstream biochemical and cellular processes that eventually lead to changes in cognition and even dementia are not well understood. Thus, a holistic or systems-level approach that aims to understand these altered processes may yield insight into new drug targets and biomarkers for AD.

Proteins are the proximate mediators of disease, integrating the effects of genetic, epigenetic, and environmental factors. It is well established that distinct mechanisms regulate expression and turnover of RNA and proteins, resulting in weak correlations in their respective levels^5^. This is mainly attributed to the complexity of the human proteome in which 90% of genes yield alternatively-spliced RNA transcripts^6,7^, and each translated protein isoform is potentially altered by up to 300 or more posttranslational modifications (PTMs)^8,9^. Protein phosphorylation is one of the key PTMs that govern signaling pathways and pathophysiological mechanisms in AD^10,11^, inspiring a large body of research that has identified several kinases including GSK-3β, CDK5, PKC, MAPK, and ROCK2 that have been implicated in the phosphorylation of tau and other key substrates in AD brain^12-16^. Conversely, reduced expression and activity of protein phosphatases like PP2A are also thought to contribute to enhanced phosphorylation of tau and other substrates^17^. Thus, quantifying total protein levels and phosphorylated proteins across AsymAD and AD stages of disease may reveal defects in kinase- or phosphatase-mediated signaling pathways and biomarkers involved in AD progression.

Advances in liquid chromatography coupled to tandem mass spectrometry (LC-MS/MS) now facilitate high-throughput detection and quantification of thousands of proteins in a given sample. Data-dependent acquisition (DDA) or shotgun approaches are the traditional methods for proteomics workflows^18^. However, a drawback of DDA approaches is that high abundance peptides are biased towards selection for tandem MS/MS and subsequent identification^19^. To improve the detection of the low abundance peptides and enhance the depth of the proteome, different off-line fractionation methods have been used to reduce the complexity of peptide mixtures in samples prior to LC-MS/MS analysis. Methods including two-dimensional gel electrophoresis^20,21^, strong cation exchange (SCX), electrostatic repulsion-hydrophilic interaction chromatography (ERLIC)^22^, and high-pH reversed-phase chromatography^23,24^ are used to increase peptide identification by separating peptides in an orthogonal dimension^25^. With the advancement of multiplex isobaric tandem mass tags (TMT), off-line fractionation and high-resolution MS, proteomic datasets are beginning to rival the depth and breadth of transcriptomic datasets^22,26^. Moreover, integrative proteomic and transcriptomic analyses in an AD post-mortem brain cohort suggests that transcriptome- and proteome-wide analyses can generate both complementary and unique information^5^. Proteomic analyses also offer the important opportunity to identify disease-specific PTMs that may participate in key pathological processes, and potentially serve as novel biomarkers and therapeutic targets.

Here we utilized a multiplex TMT MS-based proteomic approach, using similar protocols established by the Clinical Proteomics Tumor Analysis Consortium (CPTAC)^23^, to comprehensively quantify the total proteome and phosphoproteome of human post-mortem cortical cases (*n*=27) across pathology-free controls, AsymAD and symptomatic AD individuals. This led to the quantification of 11,378 unique protein groups, as well as 48,992 phosphorylated peptides representing 33,652 phosphosites. This dataset can serve as a valuable resource to help researchers elucidate the complexity of AD as it relates to proteomic signatures found in post-mortem human brain.

## Methods

### Human brain tissue

Human brain post-mortem tissues from the dorsolateral prefrontal frontal cortex (Frontal Cortex, Brodmann Area 9) were obtained from the Emory Alzheimer’s Disease Research Center (ADRC) brain bank. In total, 27 samples from 3 groups (*n*=10 control, *n*=8 AsymAD, and *n*=9 AD) were used for brain proteome and phosphoproteome analyses. Postmortem neuropathological evaluation of Aβ plaque distribution was performed according to the Consortium to Establish a Registry for Alzheimer’s Disease (CERAD) criteria^27^, while the extent of neurofibrillary tangle pathology was assessed in accordance with the Braak staging system^28^. All case metadata, including disease state, age of death, gender, sex, and apolipoprotein (*ApoE*) genotype are listed in **Supplementary Table 1 [Data citation 1]**.

### Brain tissue homogenization and protein digestion

Procedures for tissue homogenization were performed essentially as previously described^22^. Approximately 100 mg (wet tissue weight) of brain grey matter tissue was homogenized in 500 μL of 8 M urea lysis buffer (8 M urea, 100 mM Na_2_HPO_4_, pH 8.5) with HALT protease and phosphatase inhibitor cocktail (ThermoFisher) using a Bullet Blender (NextAdvance). The samples were homogenized for two full 5 min cycles at 4 °C with ∼100 μL of stainless-steel beads (0.9 to 2.0mm blend, NextAdvance). The lysates were transferred to new Eppendorf LoBind tubes followed by 3 cycles of sonication consisting of 5 s of active sonication at 30% amplitude with 15 s incubation periods on ice in between sonication pulses. Samples were then centrifuged for 5 min at 15,000 *g* and the supernatant was transferred to a new tube. Prior to further processing, protein concentration and integrity were tested by bicinchoninic acid (BCA) assay (Pierce) and SDS-PAGE, respectively. For protein digestion, 500 μg of each sample was aliquoted and volumes were normalized with additional lysis buffer. Samples were reduced with 1 mM dithiothreitol (DTT) for 30 min, followed by 5 mM iodoacetamide (IAA) alkylation in the dark for another 30 min. Samples were diluted 4-fold with 50 mM triethylammonium bicarbonate (TEAB) before incubating with Lysyl endopeptidase (Wako) at 1:100 (w/w) for 12 hr. Trypsin (Promega) was then added at a 1:50 (w/w) ratio and digestion was carried out for another 12 hr after the urea concentration was diluted to 1 M with 50 mM TEAB. The peptide solutions were desalted with a C18 Sep-Pak column (Waters). Briefly, the Sep-Pak columns were activated with 3 × 1.5 mL of methanol, then equilibrated with 6 × 1.5 mL 0.1% triflouroacetic acid (TFA). The samples were loaded after acidification to a final concentration of 1% formic acid (FA) and 0.1% TFA. Each column was washed with 6 × 1.5 mL 0.1% TFA. Elution was performed with 2 × 1.5 mL 50% acetonitrile.

### Tandem mass tag (TMT) peptide labeling

A 600 µL aliquot from each sample was pooled and the mixture was divided into 6 global internal standard (GIS) samples with a total volume of 2400 µL each, consistent with our previous work^22^, and peptide solutions were dried by vacuum (Labconco). All 27 samples from 3 groups and 6 GIS were divided and labeled using two sets of 5 mg 11 plex TMT reagents (Thermo Scientific A34808, Lot No for TMT 10-plex: SI258088, and 131C channel: SJ258847). The batch arrangement is provided in **Supplementary Table 2**. Briefly, each of the TMT reagents were dissolved in 256 μL anhydrous acetonitrile and the same channel was combined together from two 5 mg reagents. The samples were reconstituted in 400 μL of 100 mM TEAB buffer and mixed with 3.2 mg (164 μL) of the corresponding labeling reagent channel. The reaction was incubated for 1 hr and subsequently quenched with 32 μL of 5% hydroxylamine (Pierce). For each TMT plex, labeled peptides from all 11 channels were mixed and desalted with a 500 mg Sep-Pak column (Waters). The labeled peptide mixture was eluted in 4.5 mL of 50% acetonitrile and dried by vacuum.

### High-pH off-line fractionation

The off-line high pH fractionation method was adapted from the CPTAC protocol^23^. Briefly, dried samples were re-suspended in high pH loading buffer (1 mM ammonium formate, 2% (v/v) acetonitrile) and loaded onto a ZORBAX Extend 300 C18 columns (4.6 mm x 250 mm with 5 µm beads) from Agilent. An Agilent 1100 HPLC system was used to carry out the fractionation. Ammonium formate (pH 10), diluted to a concentration of 4.5 mM in 2% (v/v) acetonitrile, was used as basic-pH reverse phase solvent A, whereas 4.5 mM ammonium formate (pH 10) in 90% (v/v) acetonitrile was used as paired solvent B. The peptides were eluted in a 96-min gradient with 0%–16% mobile phase B from 7-13 min, 16%–40% B from 13-73 min, 40%–44% B from 73-77 min, 44%-60% B from 77-82 min and kept at 60% B until the end with a flow rate of 0.8 mL/min. A total of 77 individual fractions were collected across the gradient from 14 to 91 minutes, at a rate of 1 fraction per minute. The 77 individual fractions were then pooled into 24 fractions by combining 3 off-line fractions into one with alternating combinations, except for the 24th fraction, which also contained the last 5 remaining individual fractions (**Supplementary Figure 1)**. Fractions were acidified to a final concentration of 1% FA, and 5% of sample volumes were dried and reserved for proteome analysis.

### IMAC phosphorylated peptide enrichment

For phosphorylated peptide enrichment, 95% of the 24 high-pH fractions were further combined into 12 fractions in an alternating manner (1 and 13, 2 and 14, etc.). Peptide amounts were assumed to be equally distributed in all fractions. The IMAC enrichment method was performed according to CPTAC protocol with some minor modifications^23^. Briefly, 1200 μL of slurry, in which the beads/solvent ratio is 1:1 (v/v), was utilized for one batch of TMT fractions. Beads were stripped of nickel with 8 mL of 100 mM EDTA and then equilibrated with 8 mL of 50 mM FeCl_3_ both by end-to-end rotation for 30 min. To remove excess Fe^3+^ ion, beads were washed with 3 × 8 mL of water and resuspended in 2.4 mL of 1:1:1 (v/v/v) ratio of acetonitrile/methanol/0.01%(v/v) acetic acid. The beads were re-rinsed with 2.4 mL of 100% acetonitrile/0.1% TFA and divided into 12 tubes. The supernatant was removed before the peptide mixture was added. All 12 dried fractions were reconstituted in 0.4 mL of 50% acetonitrile/0.1% TFA and then diluted 1:1 with 100% acetonitrile/0.1% TFA to obtain a final 75% acetonitrile/0.1% TFA peptide solution at a concentration of 0.5 μg/μl. The peptide mixture was incubated with treated beads for 30 min with end-to-end rotation. Enriched IMAC beads were resuspended in 100 μL of 80% acetonitrile/0.1% TFA before the stage tips were conditioned. Stage tips were equilibrated with 2 × 100 μL methanol washes, 2 × 100 μL 100% acetonitrile/0.1% trifluoroacetic acid washes, followed by 2 × 100 μL of 1% FA washes. The IMAC bead slurry was loaded onto the stage tips and washed with 3 × 100 μL of 80% acetonitrile/0.1% TFA, then 3 × 100 μL of 1% FA. The phosphorylated peptides were released from IMAC beads by 3 × 100 μL 500 mM dibasic sodium phosphate (Na_2_HPO_4,_ Sigma, S9763), pH 7.0, and washed by 3 × 100 μL 1% FA. The phosphorylated peptides were eluted from stage tips by 3 × 100 μL 50% acetonitrile/0.1% FA. The phosphorylated peptide solutions were dried with vacuum.

### LC-MS/MS and TMT data acquisition on an Orbitrap Lumos mass spectrometer

Both proteome and phosphoproteome samples were run on a Fusion Lumos equipped with a NanoFlex nano-electrospray source (ThermoFisher). The same volume of loading buffer (19 μL of 0.1% TFA) was added to each of the fractions assuming equal distribution of peptide concentration across all 24 proteomic subfractions. Therefore, an equal 2 μL (1 μg equivalent) of each fraction was loaded for proteomic analysis. It was assumed that phosphorylated peptides were ∼ 1% (w/w) of all peptides. The same volume of loading buffer (7 μL, 0.1% TFA) was added to all IMAC elution samples, and of this, 2 μL (1 μg equivalent) was analyzed by mass spectrometry. All proteome and phosphoproteome samples were separated on 25 cm long 75 μm ID fused silica columns (New Objective, Woburn, MA) packed in-house with 1.9 μm Reprosil-Pur C18-AQ resin (Dr Maisch). All fractions were eluted over a 140 minute gradient using an Easy nLC 1200 (Thermofisher). The gradient started with 1% buffer B (A: water with 0.1% formic acid and B: 80% acetonitrile in water with 0.1% formic acid) and went to 7% in 3 minutes, then increased from 7% to 30% in 137 minutes, then to 95% within 5 minutes and finally staying at 95% for 25 minutes. The mass spectrometer was operated in top speed mode with 3 second cycles. Both the MS and MS/MS scans were collected in the Orbitrap. The full scan was performed with a range of 375-1500 m/z, a nominal resolution of 120,000 at 200 m/z, automatic gain control (AGC) at 4 × 10^5^, a 50 ms max injection time and an radio frequency (RF) lens setting of 30%. Higher-energy collision dissociation (HCD) MS/MS scans settings were the following: resolution of 50,000, AGC at 1 × 10^5^, isolation width of 0.7 m/z, max injection time of 105 ms, and a collision energy of 38%. Only charge states from 2+ to 7+ were chosen for tandem MS/MS. All resulting raw files (*n*=108) are provided **[Data Citation 2]**

### Protein identification and quantification

Raw data files obtained from the Orbitrap Fusion were processed using Proteome Discoverer™ (version 2.3). MS/MS spectra were searched against the UniProtKB Human proteome database (downloaded in 2015) with 90,411 total sequences with a few peptides manually added to the database, including: one APOE 2/3-specific peptide and one APOE 3/4-specific peptide which allows APOE proteotyping of samples; 4 non-tryptic C-terminal Aβ-peptides specific to Aβ38 (GAIIGLMV), Aβ40 (GAIIGLMVGGVV), Aβ42 (GAIIGLMVGGVVIA), and Aβ43 (GAIIGLMVGGVVIAT). Sequences that map to tau microtubule-binding repeat (MTBR) domains were also set as an additional entry encompassing residues 224-370 (tau 2N4R Isoform, 1-441) in the Uniprot sequence, while all tau isoform sequences were modified by removing MTBR peptides and replicated as new “deltaMTBR” entries^29,30^. Separation and quantification of these peptide sequences facilitated the investigation of APOE allele, Aβ peptide and MTBR peptide-specific regulation of biology in AD datasets^29^ **[Data Citation 3]**. The respective FASTA database used in this study was deposited on Synapse (syn20820455). The SEQUEST HT search engine was used and parameters were identical for both total and IMAC proteomes and specified as the following: fully-tryptic specificity; maximum of two missed cleavages; minimum peptide length of 6; fixed modifications for TMT tags on lysine residues and peptide N-termini (+ 229.162932 Da) and carbamidomethylation of cysteine residues (+ 57.02146 Da); variable modifications for oxidation of methionine residues (+ 15.99492 Da); deamidation of asparagine and glutamine (+ 0.984 Da); phosphorylation of serine, threonine and tyrosine (+ 79.9663 Da); precursor mass tolerance of 20 ppm; fragment mass tolerance of 0.05 daltons. Percolator was used to filter peptide spectral matches (PSM) and peptides to a false discovery rate (FDR) of less than 1% using target-decoy strategy. The phosphosite localization site threshold was set to 0.75, ensuring <5% false-localization rate (FLR) of PTM assignments as described^31^. Following spectral assignment, peptides were assembled into proteins and were further filtered based on the combined probabilities of their constituent peptides to a final FDR of 1%. In cases of redundancy, shared peptides were assigned to the protein sequence in adherence with the principles of parsimony. Reporter ions were quantified from MS2 scans using an integration tolerance of 20 ppm with the most confident centroid setting. The search results and TMT quantification are included **[Data Citation 4]**.

### Data Records

All files have been deposited on Synapse (<u>syn20820053</u>).These include sample trait (<u>syn20820456</u>, **Data Citation 1**), all mass spectrometry raw files (*n*=108) from both total proteome and phosphoproteome (<u>syn20820454</u> **Data Citation 2**), the FASTA database (<u>syn20820455</u>, **Data Citation 3**), search result (<u>syn20820472</u>, **Data Citation 4**), and the ANOVA analysis input and result (<u>syn21986491</u>, **Data Citation 5**).

### Technical Validation

#### Deep dive proteome of human AD brain

We utilized a modified version of the CPTAC protocol to identify the total proteome and phosphoproteome from the same cases across different stages of AD. Control, AsymAD and AD tissues were randomized across the 3 batches (each containing 11 samples) with 9 individual cases per batch (**Figure 1a** and **Supplementary tables 1** and **2**). Two TMT channels in each batch were dedicated to global reference internal standards (GIS), representing an equivalent amount of pooled peptides from all cases, which allows assessment of the intra- and inter-batch variance^22^. To reduce sample complexity and increase proteome depth prior to LC-MS/MS, we employed off-line high-pH reversed-phase fractionation essentially as described in the CPTAC protocol^23^. A total of 77 individual fractions were collected and combined into 24 fractions for total proteome analysis (**Supplementary Figure 1**) for each batch. A step-wise concatenation strategy was used for pooling the fractions. A total of 5% of the material by volume was used for total proteome analysis, and the remaining 95% of the sample was used for phosphopeptide enrichment by immobilized affinity chromatography (IMAC) with Fe^3+^-loaded nitrilotriacetic acid (NTA) beads. Each of the 24 fractions were pooled into 12 subfractions prior to IMAC. Both the total proteome (*n*=72 fractions across 3 batches) and phosphoproteome (*n*=36 fractions across 3 batches) were analyzed by LC-MS/MS with high-resolution precursor and MS/MS scans on an Orbitrap Fusion Lumos mass spectrometer.

**Figure 1.**
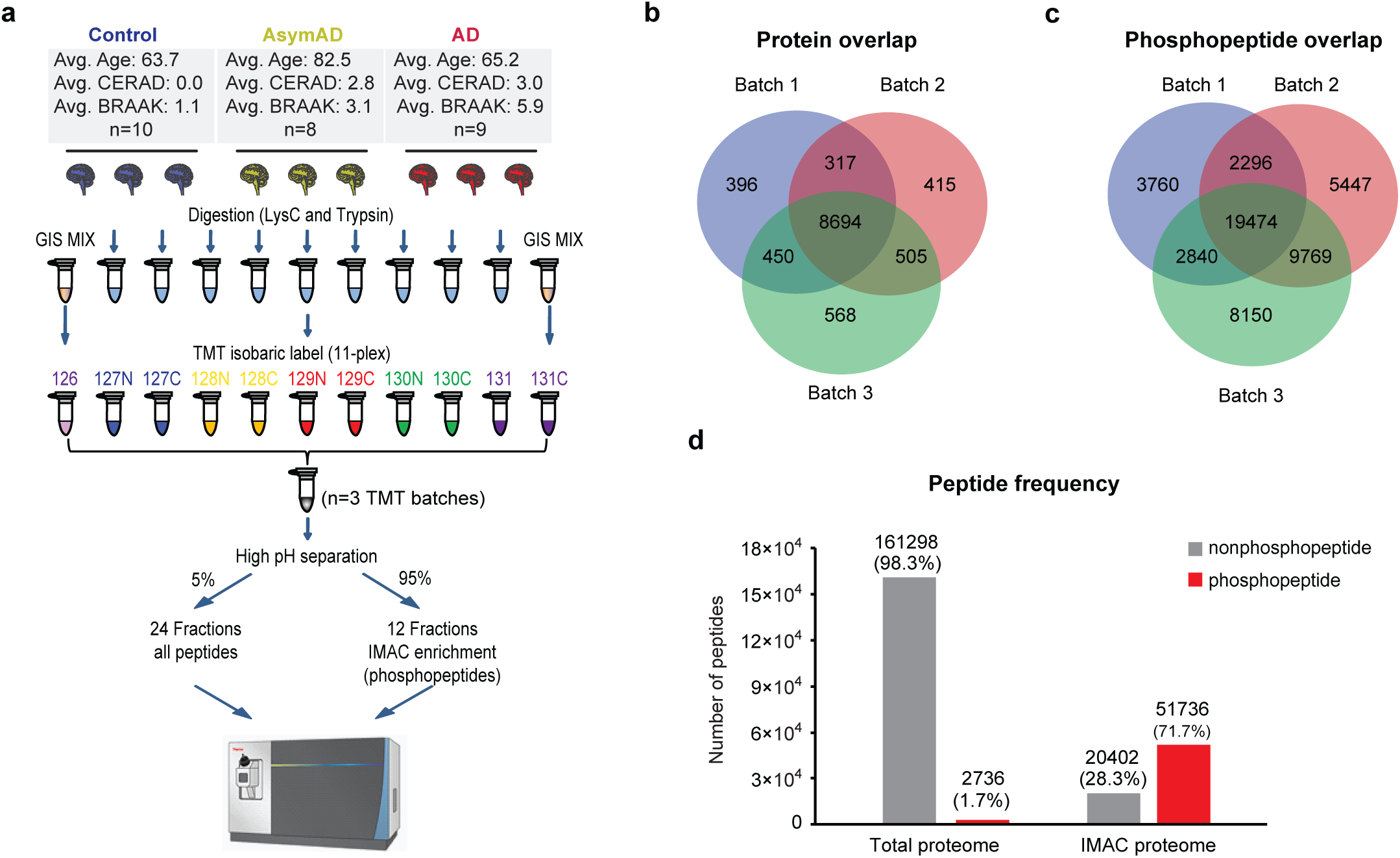
Deep coverage of human brain proteome and phosphoproteome in Alzheimer’s disease using TMT. (a) TMT Workflow. There were 27 human post-mortem tissues from the dorsolateral prefrontal frontal cortex, which were digested and labeled with 3 batches of 11 plex TMT reagents. Two pooled global internal standards (GIS) were labeled with channels 126 and 131C in each batch. There were 9 samples from 3 groups (control, AsymAD and AD) randomized and labeled in the remainder channels. The labeled sample mixture was fractionated by off-line high-pH reversed-phase chromatography. Whole proteome analysis was conducted using 5% of peptides from each fraction. Phosphopeptides were enriched from 95% of the samples by immobilized affinity chromatography (IMAC) with Fe^3+^-loaded nitrilotriacetic acid (NTA) beads. All fractions were analyzed on an Orbitrap Fusion Lumos Tribrid mass spectrometer. (b) Venn diagram of total proteome proteins from 3 batches. Approximately 10,000 protein groups were identified from each batch, with 8,694 shared protein groups found in all 3 batches, and 9,966 protein groups found in at least 2 batches. (c) Venn diagram of phosphopeptides identified in 3 batches. There were 19,474 shared phosphopeptides found across all 3 batches. (d) Peptide composition from both total proteome and IMAC proteome were illustrated. Non-phosphopeptides (*grey*) and phosphopeptides (*red*) are shown. Phosphopeptides were greatly enriched in IMAC proteome (71%), compared with the total proteome (1.7%).

For the total proteome runs, a total of 164,034 unique peptides were identified that mapped to 11,378 protein groups at a 1% FDR on the peptide spectrum match (PSM) level across all batches, which represented 10,373 coding gene products. The total numbers of identified peptides, proteins and PSMs for all batches in the total proteome are listed in **Table 1**. For each batch, there were approximately 10,000 protein groups identified (**Figure 1b**), which was comparable to the depth achieved in the CPTAC protocol using different tissue sources^32,33^. The confidence of identification for peptide and protein is highly related with the number of PSMs and unique peptides. In the total proteome dataset, more than 77% of the proteins were identified with 2 or more unique peptides (**Figure 2a**), while each unique peptide averaged approximately 6 PSMs (**Table 1**). Approximately 93% of all proteins were identified with at least 2 PSMs (**Figure 2b**).

**Table 1.** Protein and peptide identification from total proteome.

| Batch Number | Protein Groups | Unique Peptides | PSM | Phosphopeptides<br>(no enrichment) |
| --- | --- | --- | --- | --- |
| 1 | 9,857 | 112,825 | 289,129 | 1,719 |
| 2 | 9,931 | 116,558 | 315,565 | 1,785 |
| 3 | 10,217 | 127,056 | 351,413 | 1,809 |
| Total | 11,378 | 164,034 | 956,107 | 2,758 |

**Figure 2.**
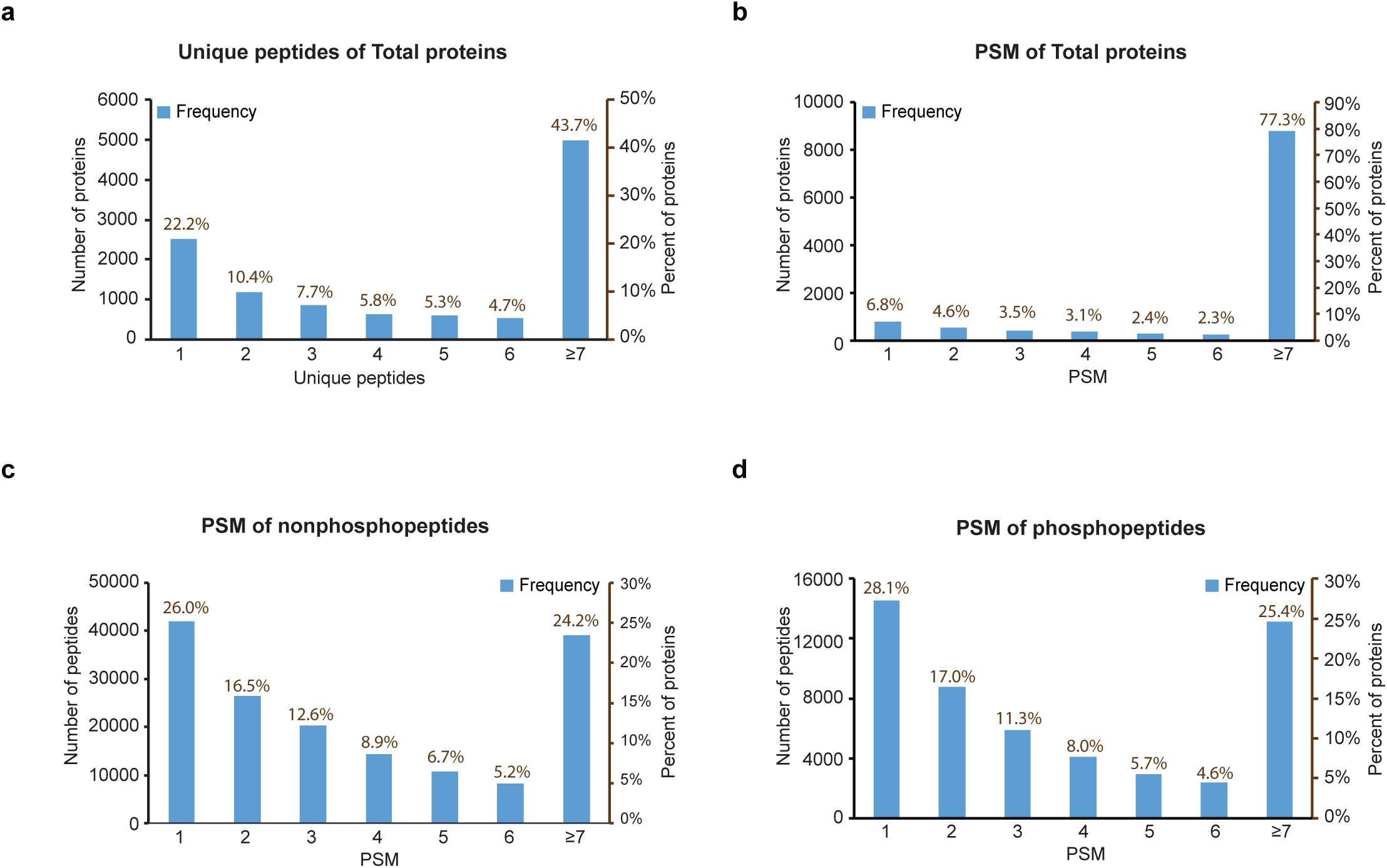
Assessment of protein and peptide identification by peptide spectral matches for both proteome and phosphoproteome. (a) The frequency of protein groups (*blue bars*) per unique peptide bin for the total proteome. The percent of proteins per each unique peptide of all protein groups was also expressed. Approximately 77% of all protein groups identified were matched by at least 2 unique peptides. (b) The frequency of protein groups (*blue bars*) per peptide spectral match (PSM) bin for the whole proteome. The percent of proteins of total protein groups identified per each PSM bin was also quantified. More than 93% of the proteins identified were sampled by 2 or more PSMs. (c) The frequency of non-phosphopeptides (*blue bars*) matching to different PSM bins from total proteome. The percentage of peptides per each PSM bin out of all non-phosphopeptides identified was also quantified. There were 119,343 non-phosphopeptides identified with 2 or more PSMs, approximately 74% of all non-phosphopeptides identified. (d) The frequency of phosphopeptides (*blue bars*) matching to different PSM bins from the IMAC-enriched phosphoproteome. The percentage of peptides per each PSM bin out of all phosphopeptides identified from the IMAC-enriched proteome was also quantified. There were 37,195 phosphopeptides identified with 2 or more PSMs, making up approximately 72% of all IMAC phosphopeptides.

To obtain deep coverage of the phosphoproteome, enrichment strategies are usually applied due to the relatively low abundance of phosphorylation. To assess the quality of our IMAC phosphopeptide enrichment method, we calculated the percent phosphopeptide content (peptide level) in both the total proteome and IMAC phosphoproteome datasets (**Tables 1** and **2**, and **Figure 1d**). The total proteome identified a total of 164,034 unique peptides, and approximately 2% were phosphopeptides (**Figure 1d**). Although the IMAC dataset identified less peptides overall (*n*=72,138), approximately 71% of the IMAC proteome were phosphopeptides (**Table 2**, and **Figure 1d**). The IMAC enrichment method therefore led to an 18-fold increase in phosphopeptide identification using half of the instrument time. We set the threshold of phosphosite identification was set to 0.75 by SEQUEST, estimating less than 5% false localization rate (FLR) of each assigned site. After filtering, approximately 83% of all phosphosites identified had localization scores greater than or equal to 0.99. There were 51,736 phosphorylated peptides with 33,652 unique individual phosphosites in total mapping to 8,415 proteins. A total of 34,379 of the phosphorylated peptides were identified in at least two of the three IMAC TMT batches (**Figure 1c**). These figures are similar to the depth reported using the same protocol from breast cancer tissue^34^. The numbers of identified peptides in each IMAC batch are listed in **Table 2** with calculation of phosphorylation enrichment at both the total peptide and PSM level. Of note, the phosphorylated peptides showed slightly higher level of enrichment at the PSM level (83.55% as average) than peptide level (71.72% as average), which indicates that phosphopeptides were more intense and thus more frequently sequenced by LC-MS/MS than an average unmodified peptide in the phosphoproteome. Indeed, each non-phosphopeptide was identified by an average of 3.5 PSMs, whereas each phosphopeptide was identified by an average of 7 PSMs (**Table 2**). In total, 74% of all phosphopeptides identified by IMAC enrichment had two or more PSMs, which is consistent with the frequency of PSMs for peptides identified from the total proteome (**Figure 2c and 2d**). Thus, although phosphopeptides were highly-enriched in the IMAC proteome, they were sampled at a rate generally consistent with non-phosphopeptides from the total proteome, allowing greater sequencing depth of the phosphoproteome.

**Table 2.** Peptide identification from IMAC samples.

| Batch Number | Total Peptide | Phospho-peptide | % of Enrichment (Peptide level) | Total PSM | Phospho-PSM | % of Enrichment (PSM level) |
| --- | --- | --- | --- | --- | --- | --- |
| 1 | 34,056 | 28,370 | 83.30% | 113,894 | 102,905 | 90.35% |
| 2 | 53,349 | 36,986 | 69.33% | 157,279 | 118,150 | 75.12% |
| 3 | 50,922 | 40,233 | 79.01% | 163,477 | 142,074 | 86.91% |
| Total | 72,138 | 51,736 | 71.72% | 434,650 | 363,129 | 83.55% |

### Assessing intra- and inter batch variance utilizing a pooled global internal standard

A major advantage of TMT approaches is the ability to quantify multiple samples in a single run, thereby critically reducing overall MS instrument time. This becomes especially important when the total number increases to dozens or even hundreds of samples^30,35,36^. Typically, one or more TMT channels are dedicated for global internal standard(s) (GIS) and included in all batches, which can be used to normalize the measurement for protein or peptide signal from all samples across all batches*22*. In this study, we included two pooled reference standards in each 11-plex TMT batch (channels 126 and 131C), which allows normalization within and across TMT batches (**Figure 1a, Supplementary Tables 1** and **2, data citation 1**). The two reference standards essentially serve as technical replicates (i.e., a null-experiment), which can be used to assess the variance in measurements. Thus, the degree of the signal variation between two internal GIS channels can be used as a threshold to further filter out the poor quantitation data. Indeed, the signal of proteins and peptides from 126 and 131C channels were very consistent and showed very good linear correlation across all three batches (**Figure 3a**). We also consistently observed a strong correlation at both the peptide level from the total proteome (**Figure 3b**) and phosphoproteome (**Figure 3c**). Notably, some peptides exhibited large variation in signal between the two pooled standard channels, especially those peptides with lower total signal abundance as previously described^22^. According to the central limit theorem, the log_2_ ratio for the two GIS channels (log_2_ TMT channels 126/131C) should fit a standard Gaussian distribution with the mean at or near zero (**Supplemental Figure 2a**), which can be used to assess the technical variation of measurements^37,38^. This allows end-users of the datasets to impose a filtering criterion that can be used to remove peptides or proteins that do not meet variance metrics (>2 standard deviations (SD) from the mean). Following this filtering criteria, a total of 1,123 peptides were filtered out of the analyses due to large variance, equivalent to ∼4% of all quantitated peptides in IMAC Batch 1. In batch 2 and batch 3, there were 2,465 and 2,289 peptides filtered out by the >2SD standard, representing 5% of all peptides identified respectively (**Supplementary Figure 2b**).

**Figure 3.**
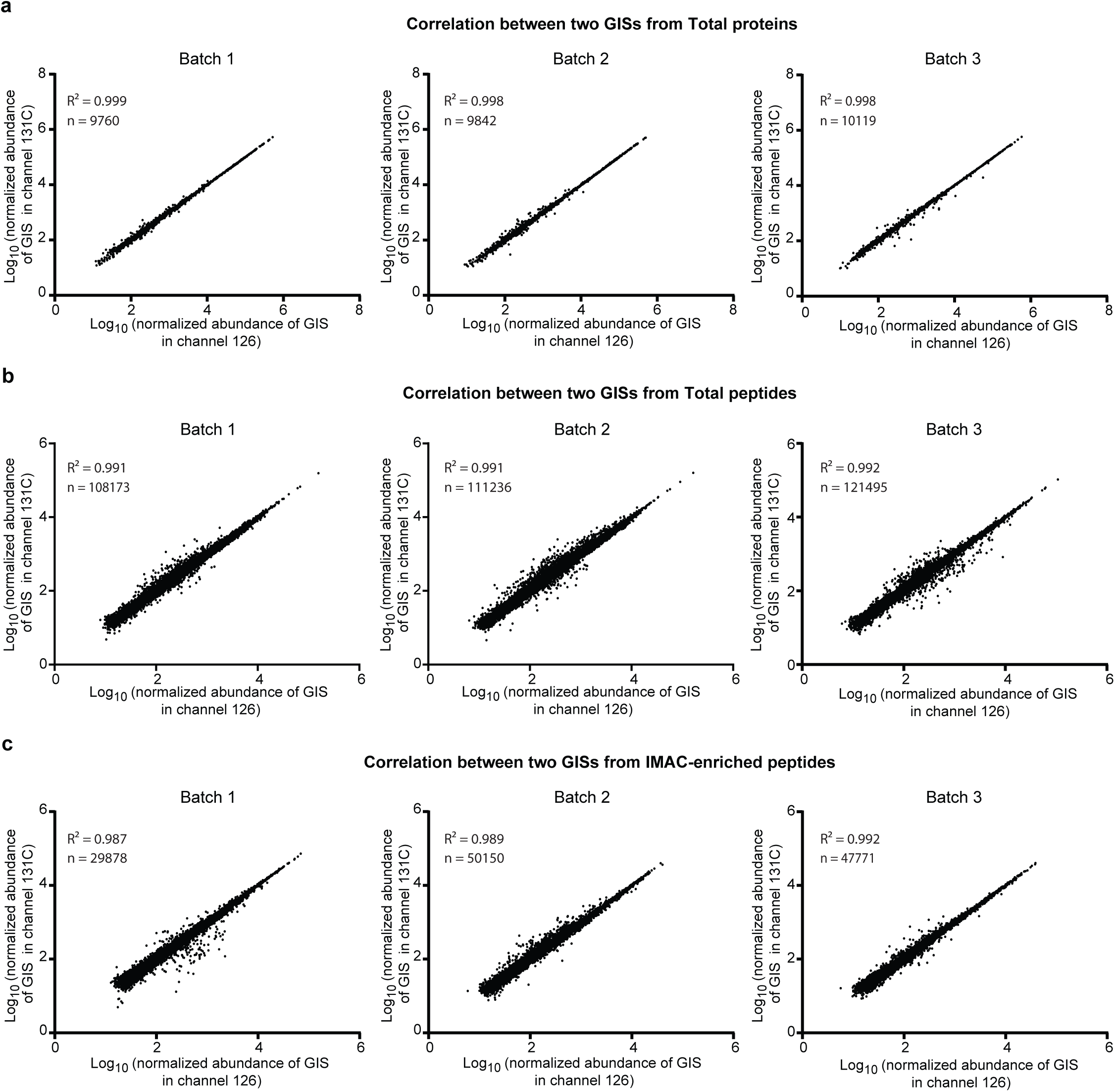
Correlation of two pooled GIS samples across proteome and phosphoproteome batches. (a) Correlation of two GIS samples, which were in the TMT channels 126 and 131C, in 3 total proteome batches at the protein level. Strong correlation was observed between 2 GIS channels in all 3 batches. (b) Correlation of 2 GIS samples in 3 whole proteome batches at the peptide level. (c) Correlation of 2 GIS samples in 3 IMAC-enriched phosphoproteome batches at the peptide level. Normalized 126 and 131C channel intensities illustrate tight correlation.

It is also worth noting that batch effects may be significant when the sample number is large and the variation due to sample preparation cannot be ignored. In this case, post-hoc data normalization strategies should be employed to remove these batch effects^39,40^. In this project, however, this step was not necessary given the relatively modest sample size (*n*=27) and since all samples were digested at the same time.

### Assessing amyloid levels and tau phosphorylation

Aβ plaque and hyperphosphorylated tau neurofibrillary tangle (NFT) accumulation in the brain are the core pathological hallmarks of AD^41,42^. Thus, as a quality control of our measurements, we assessed the levels of Aβ and tau in our dataset. To confirm increased Aβ levels in diseased cases, the ion intensities from first two tryptic peptides of Aβ were used as a surrogate for amyloid levels in the brain^5^, corresponding to residues 6-16 (Peptide 1) and 17-28 (Peptide 2) of the Aβ sequence, since the C-terminal non-tryptic peptides were not stably detected in all batches. Indeed, both of these two peptides showed significant increase in AsymAD and AD groups comparing with control samples (**Figure 4a, Data Citation 5**). Additionally, measurements of Peptide 1 and Peptide 2 were highly correlated (**Figure 4b**). Given this, the sum intensity of the two peptides was used to represent Aβ levels in each sample, which showed significant increase in both AsymAD and AD samples when compared to Control (**Figure 4c**).

**Figure 4.**
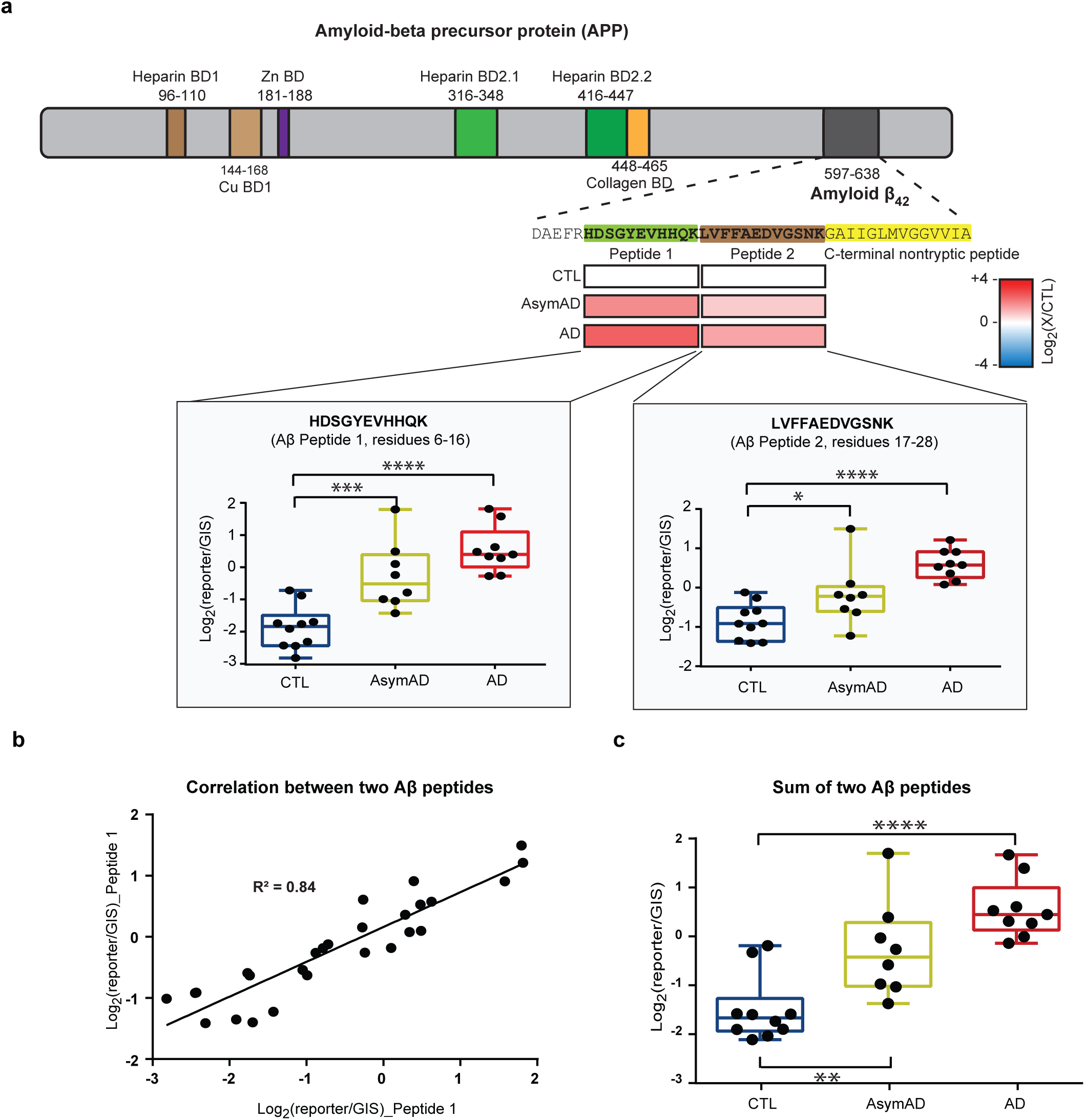
Quantification of Aβ peptides. (a) Amyloid-beta precursor protein (APP) protein domain map showing the Aβ_42_ region (*dark grey*). The abundances of tryptic Aβ peptide 1 (residue 6-16, *green*) and peptide 2 (residue 17-28, *brown*) was normalized to global internal reference standards (GIS) and quantified in control (CTL), AsymAD, and AD cases. The log_2_ abundances of diseased cases compared with controls of Peptide 1 and Peptide 2 were illustrated as increased [Log_2_(X/CTL) = +4, *red*] or decreased [Log_2_(X/CTL) = -4, *blue*]. Both Peptide 1 and Peptide 2 were significantly increased in AD and AsymAD groups. (b) Correlation of the abundances of Peptide 1 and Peptide 2 from the same samples. (c) The GIS-normalized abundances of the sum of two Aβ peptides in CTL, AsymAD, and AD cases. The summed level of Aβ peptides was significantly changed between control and AD groups, with both AD groups (****, p<0.0001 one-way ANOVA) and AsymAD groups (**, p<0.01) containing significantly-increased Aβ.

Another hallmark of AD is hyperphosphorylated tau^43^, which is the core component of neurofibrillary tangles (NFTs) in diseased neurons. Remarkably, 21 phosphorylated tau peptides were detected in total proteome even without IMAC enrichment, which highlights the robust phosphorylation of this protein in AD brain (**Figure 5a**). After IMAC enrichment, a total of 112 tau phosphopeptides were identified. Of note, there were 47 peptides containing two or more phosphosites, which was close to 42% of all phosphopeptides mapped to tau **(Figure 5a)**. Since the MTBR domains form the core of neurofibrillary tangles and is required to seed tau aggregation^44,45^, it was set as an additional protein entry as MAPT-MTBR within the database, while all other tau isoforms were replaced as new “deltaMTBR” entries after the MTBR sequence was removed from the original sequences. As shown in **Figure 5b and 5c**, both MAPTdeltaMTBR (MAPTΔMTBR) and MAPT-MTBR show differences between AD and control groups. However, as expected, the effect size (log_2_ fold change) for the tau MTBR is larger than the tau with the ΔMTBR in AD. The tau MTBR sequenced from the phosphoproteome, which contained stronger phosphopeptides signal, yielded even better separation between AD and control groups compared with MTBR sequenced from the total proteome.

**Figure 5.**
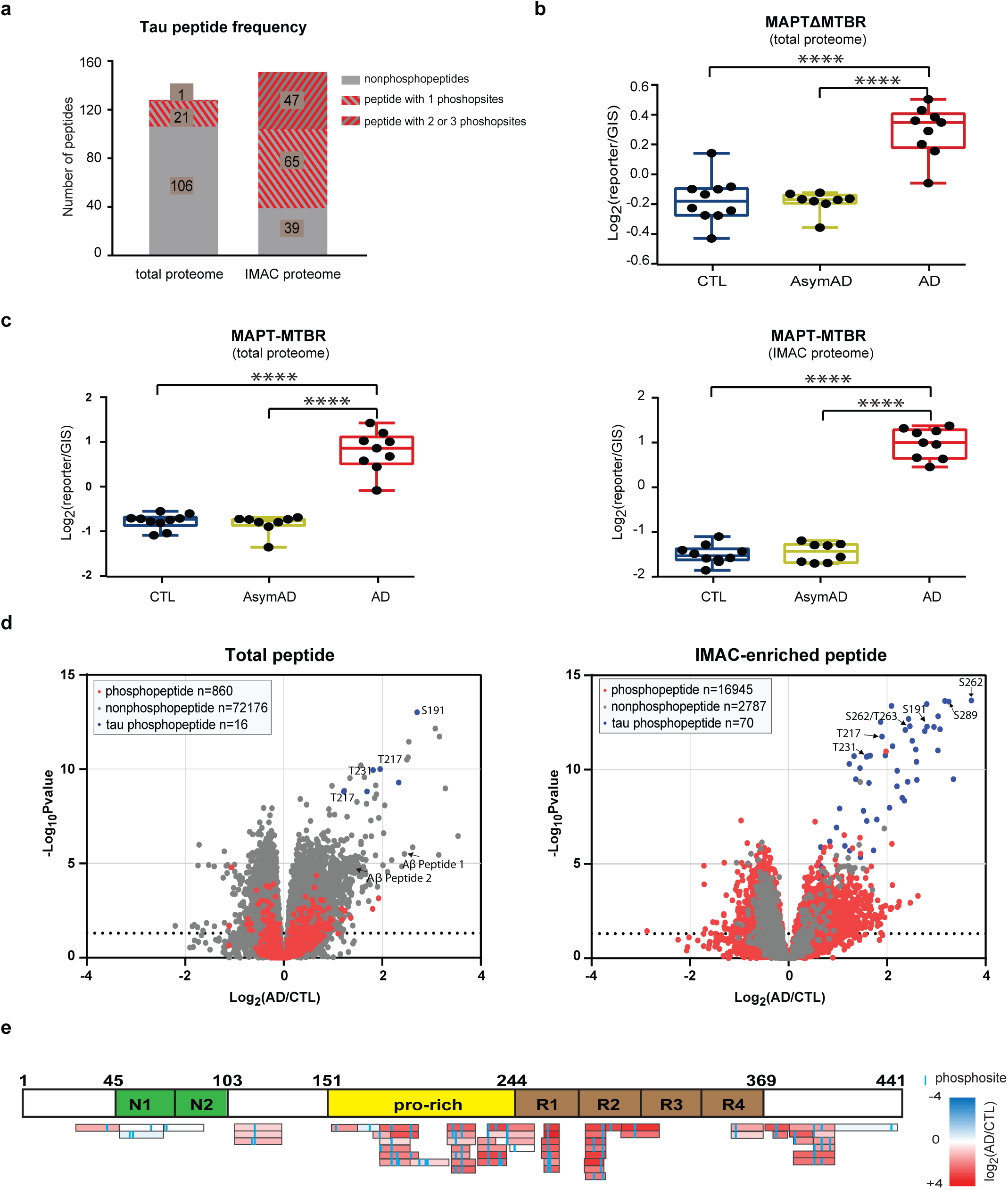
Tau phosphopeptides are highly enriched in AD. (a) Tau peptide composition from both total proteome and IMAC proteome. In the total proteome, 17% of peptides mapping to Tau were phosphopeptides, while 74% of all Tau peptides in the IMAC proteome were phosphopeptides. Approximately 42% of the Tau phosphopeptides in the IMAC proteome contained 2 or more phosphosites. (b) The GIS-normalized abundances of MAPTΔMTBR in CTL, AsymAD, and AD cases. The MTBR was removed from MAPT and set as an additional sequence in the FASTA. AD groups showed significant increase compared to control and AsymAD groups. (c) The GIS-normalized abundances of MAPT-MTBR in CTL, AsymAD, and AD cases. AD groups showed significant increase comparing to control and AsymAD groups. Compared to MAPTΔMTBR, MTBR domains showed better separation from both the total proteome and phosphoproteome data. (d) Volcano plots showing log_2_ abundance fold changes (AD/CTL) of non-phosphopeptides (*grey*) and phosphopeptides (*red*) and log_10_-transformed p-values after one-way ANOVA across three groups (CTL/AsymAD/AD) in the total proteome dataset (*left*) and IMAC-enriched phosphoproteome dataset (*right*). In order to achieve better accuracy, only the peptides quantified across all 3 batches were plotted. Several phosphorylated MAPT peptides (*blue*) identified from both total proteome and IMAC proteome were significantly increased in AD compared with CTL and labeled. Both tryptic Aβ peptides increased in the AD group. (e) The log_2_ fold changes in AD over CTL brains of tau phosphopeptides were largely increased in the Proline-rich (Pro-rich) domain (yellow) and MTBR domains (brown). Unchanged phosphopeptides also mapped to the N-terminal acidic domains (green). The log_2_ abundances of Tau peptides quantified in at least two batches were illustrated ranging from increased [Log2(X/CTL) = +4, *red*] to decreased [Log2(X/CTL) = -4, *blue*].

A one-way ANOVA of peptide levels across three groups (CTL, AsymAD and AD) was also performed (**Data Citation 5**), and peptide volcano plots were calculated, showing log_2_ fold changes and log_10_-transformed p-values of peptides from both the total proteome and phosphoproteome datasets (**Figure 5d**). In total proteome peptide data, we observed both Peptide 1 and Peptide 2 from Aβ to be significantly increased in AD when compared to controls. Additionally, the tau phosphopeptides were among the most changed peptides between AD and control, with significantly increased peptide abundances (**Figure 5d**). In agreement with this, tau phosphopeptides were among the most significantly-changed peptides in the IMAC proteome as well. To illustrate this, all tau phosphopeptides quantified in more than two batches were colored according to the degree of fold change between AD and control from IMAC proteome (**Figure 5e**). Importantly, the IMAC enrichment allowed deep sequencing and quantification of phosphopeptides mapping to the Proline-rich (Pro-rich) domain (residues 103-244) and MTBR (residues 244-368) domain^46^. Both these regions showed the most consistently increasing in abundance in AD compared with other regions of the tau protein.

### Usage Notes

Ultimately, these deep human brain proteomic and phosphoproteomic datasets serve as a valuable resource for a variety of research endeavors including, but not limited to, the following applications:

### Use case 1: Protein abundance at steady state

This dataset provides a reference for relative protein abundance in brain, especially if an investigator wants to determine whether their protein of interest is abundantly expressed in human brain. [**Data Citation 4**].

### Use case 2: AD stage-specific differential protein expression

There were three separate clinical and pathological groups of human post-mortem tissues representing three stages of AD. One can compare the expression differences between different stages at the protein, peptide or phosphopeptide level. The volcano plots shown in **Figure 5d** displays the substantive changes in peptide levels between AD and control groups. The same analysis between AsymAD and control can also be applied. This analysis also includes the quantification of peptides with and without phosphorylation sites in the same peptide within the same sample, which can greatly benefit the investigators working to fully describe the phosphorylation stoichiometry of certain proteins [**Data Citation 4**].

### Use case 3: Protein co-expression network analysis

In this dataset, there were more than 10,000 proteins quantified which is more than enough to conduct systems-level analysis. WeiGhted Co-expression Network Analysis (WGCNA) and related algorithms can be utilized for systems-based network analyses, which generate modules of proteins clustered by correlated expression patterns^5,30,36^. The protein clusters can then be correlated to molecular functions and pathways. These programs can also be used to correlate expression clusters to various biological traits. Furthermore, the cell-type specificity of individual proteins may be investigated according to the module membership of a protein and the brain cell-type enrichment data for that particular module.

### Use Case 4: Identification and quantification of signaling pathways

Pathway analysis is routine with software^47^ or web services^48,49^ to analyze different high-throughput omics data, like genomics, transcriptomics, proteomics, lipidomics and metabolomics. Pathway analyses help to organize a list of proteins into a cohesive list of pathway maps to interpret proteomics results. These analyses have proved to be a very powerful interpretation tool in biological research, facilitating novel insights in disparate fields including development^50^, apoptosis^51^, cancer^52,53^, and other diseases^54,55^. Several biological pathways have been linked to AD using similar methods^30,56-58^. Given the excellent coverage of the AD proteome and phosphoproteme from the same samples described here, this dataset may therefore serve as a useful resource for pathway analysis.

### Use Case 5: Domain or motif analysis

A protein domain or motif is a part of a given protein sequence that serves as a substrate for kinases or other enzymes to recognize and chemically modify, and is replicated in other sequences in the proteome, playing conserved roles in protein function^59,60^. Recent advances in genomics and proteomics sequencing following the development of bioinformatics^61,62^ making possible large-scale domain or motif analyses. As kinases reliably phosphorylate motif sequences specific to that enzyme, the altered phosphorylation of certain motifs may reflect impaired kinase dynamics in AD. Given the enhanced coverage of the AD proteome and phosphoproteome, this dataset can be an excellent tool for AD-related domain or motif analysis.

### Use case 6: Targeted proteomics

Due to the multiplexing nature of the TMT method, proteomic sample processing has become increasingly high throughput and a more popular mode of research. As innovative technical advances in instrumentation, computing and processing have steadily improved, TMT-labeled peptide analyses have begun to be applied to targeted proteomic methods, such as TOMAHAQ^63^. In this dataset, we have identified 164,034 peptides and 51,736 phosphopeptides through TMT isobaric labeling. Importantly, this dataset includes peptide-specific characteristics such as intensity, charge and modification state, which can serve as a resource to reference for targeted proteome analyses in the future.

### Data Citations

1. Ping, L, Duong D. M., & Seyfried, NT SAGE BIONETWORKS Synapse: syn20820456 (2019)
2. Ping, L, Duong D. M., & Seyfried, NT SAGE BIONETWORKS Synapse: syn20820454 (2019)
3. Ping, L, Duong D. M., & Seyfried, NT SAGE BIONETWORKS Synapse: syn20820455 (2019)
4. Ping, L, Duong D. M., & Seyfried, NT SAGE BIONETWORKS Synapse: syn20820472 (2019)
5. Ping, L, Duong D. M., & Seyfried, NT SAGE BIONETWORKS Synapse: syn21986491 (2020)

### Code Availability

The authors do not have code specific to this work to disclose.

## Acknowledgments

We are grateful to the patients and families that donate tissue samples to the Emory University brain bank and for their contributions to this study. Support for this research was provided by funding from the National Institute on Aging (R01AG053960, R01AG061800, RF1AG057471, RF1AG057470, R01AG057339, RF1AG062181), the Accelerating Medicine Partnership for AD (U01AG046161 and U01AG061357) and the Emory Alzheimer’s Disease Research Center (P50AG025688). All raw data, case traits and analyses related to this manuscript are available at https://www.synapse.org/#!Synapse:syn20820053

## Author contributions

Conceptualization, L.P., D.M.D., J.J.L., A.I.L., N.T.S.; Methodology, L.P., D.M.D., L.Y., M.G., N.T.S.; Investigation and Formal Analysis, L.P., D.M.D., S.R.K., N.T.S.; Writing – Original Draft, L.P., S.R.K., N.T.S.; Writing – Review and Editing, L.P., S.R.K., D.M.D., J.J.L., A.I.L., N.T.S.; Funding Acquisition, A.I.L. and N.T.S.; Supervision, N.T.S.

## Supporting Information

**Supplementary Figure 1.**
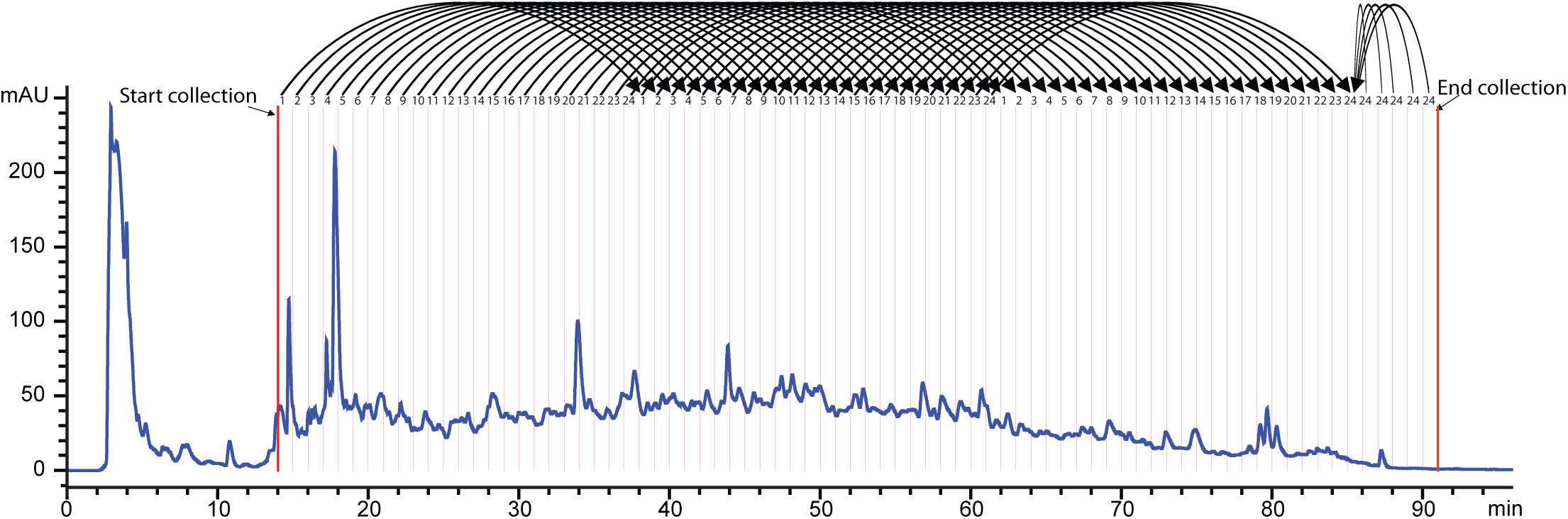
Chromatography of off-line high-pH peptide separation. The TMT-labeled peptide mixture was separated on a 96 min gradient. In total, 77 fractions were collected from 14 to 91 minutes, with 1 fraction collected every min, which were further combined into 24 fractions. The 24 fractions were further combined into 12 fractions in an alternating manner (1 and 13, 2 and 14, etc.) for phosphoproteome analysis.

**Supplementary Figure 2.**
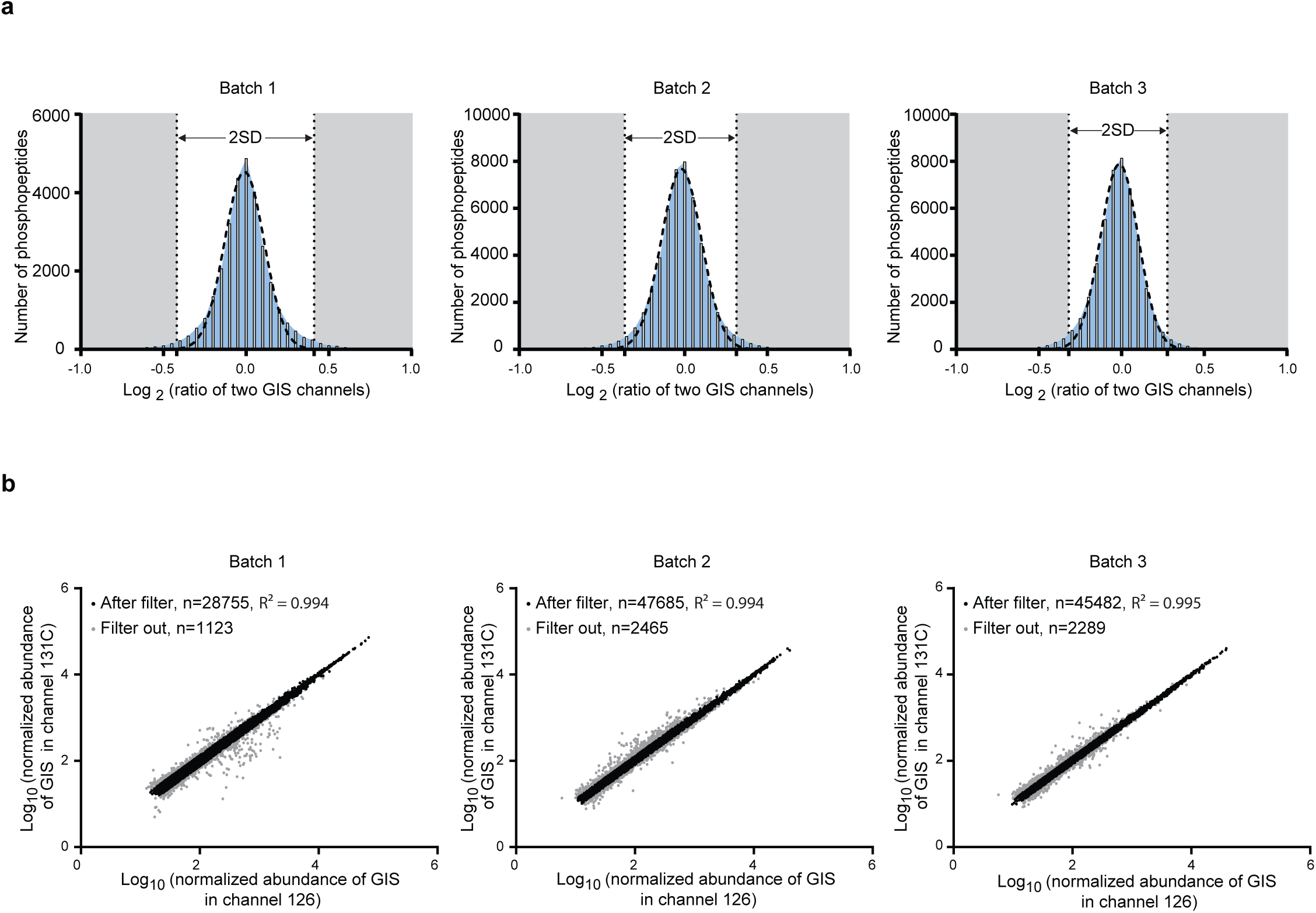
Evaluation of stability of quantification using 2 GIS samples. (a) The Gaussian distribution of log_2_ ratios of the normalized abundance of 126/131C. The Gaussian curves were fit for all 3 batches using Graphpad Prism6. The peptides with log_2_(126/131C) ratios >2 standard deviation from 0 are highlighted (*grey*). (b) The adjusted correlation of normalized abundance from 126 and 131 channels after log_2_ ratio of (normalized abundance of 126/131) data were filtered with 2 standard deviations. Those peptides that were >2 standard deviation (*grey points*) were excluded from further analyses, while peptides with ≤2 SD were included (*black dots*). After filtering out variable peptides from the IMAC proteome, the GIS channels were better correlated.

